# Genomic evidence for a chemical link between redox conditions and microbial community composition

**DOI:** 10.1101/2021.05.31.446500

**Authors:** Jeffrey M. Dick, Jingqiang Tan

## Abstract

Environmental influences on community structure are often assessed through multivariate analyses in order to relate microbial abundances to separately measured physicochemical variables. However, genes and proteins are themselves chemical entities; in combination with genome databases, differences in microbial abundances directly encode for chemical variability. We predicted that the carbon oxidation state of inferred community proteomes, obtained by combining taxonomic abundances from published 16S rRNA gene sequencing datasets with predicted microbial proteomes from the NCBI Reference Sequence (RefSeq) database, would reflect environmental oxidation-reduction conditions in various natural and engineered settings including shale gas wells. Our analysis confirms the geobiochemical predictions for environmental redox gradients within and between hydrothermal systems and stratified lakes and marine environments. Where they are present, a common set of taxonomic groups (Gamma- and Deltaproteobacteria and Clostridia) act as drivers of the community-level differences in oxidation state, whereas Flavobacteria most often oppose the overall changes. The geobiochemical signal is largest for the steep redox gradients associated with hydrothermal systems and between surface water and produced fluids from shale gas wells, demonstrating the ability to determine the magnitude of redox effects on microbial communities from 16S sequencing alone.

## Introduction

How environment affects community structure is a major question for microbial ecology. Physicochemical conditions affect the energetics of metabolic reactions (e.g. ***Lin et al., 2016***) and consequently the distribution of organisms with specific metabolic capabilities. In some environments such as sediments, there are well-known links between inorganic redox couples and metabolic types. However, at a global scale, salinity is thought to be the major driver of community structure (***Lozupone and Knight, 2007***; ***Thompson et al., 2017***).

The elucidation of environmental drivers of community structure depends on two types of data: environmental DNA sequencing (e.g. 16S amplicon or shotgun metagenomes) and physicochemical measurements. Many studies have used multivariate techniques, particularly ordination, to relate microbial abundances determined from sequence data to measured physicochemical variables (***Ramette, 2007***). Despite the sophistication of these techniques, there are some drawbacks: the environmental factors are represented by dataset-dependent synthetic variables that can be difficult to interpret (***Ramette, 2007***), and the analyses are inherently correlative (***James and McCulloch, 1990***). A complementary approach would be to use the particular biomolecular compositions of different organisms to derive chemical metrics, which have a stable definition that enables comparisons across datasets. Such a chemical representation is also a precursor to thermodynamic models that yield quantitative predictions of the effects of the physicochemical environment on the relative abundances of different microbial organisms (***Dick and Shock, 2013***).

The analysis of environmental DNA sequences is most often regarded as a taxonomic classification problem: What organisms are there? However, from a geobiochemical perspective, microbial ecosystems can also be viewed as chemical systems, which prompts a different question: What are the organisms made of? One approach to answer this question is to obtain the elemental compositions of proteins coded by the genomes of different organisms. The methodological advance in this study is to combine proteins coded by whole-genome reference sequences with taxonomic profiles from 16S rRNA gene sequences in order to infer community proteomes. These inferred proteomes have a definite elemental composition and therefore provide a novel source of information about biochemical differences between microbial communities.

Our guiding hypothesis is that physicochemical gradients shape the chemical composition of microbial communities. The specific predictions we make are that where an environment is more reducing, the community proteomes would have a lower average oxidation state of carbon (*Z*_C_), and where an environment is more saline (which is a condition associated with lower water activity), the community proteomes would have a lower stoichiometric hydration state 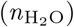. This choice of variables reflects the anticipation that oxidation-reduction and hydration-dehydration reactions are major types of biochemical transformations in metabolism and, by extension, at the ecosystem scale. We previously described the rationale and derivation of *Z*_C_ and 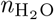 applied to shotgun metagenomic data for redox gradients and salinity gradients (***Dick et al., 2019, 2020***). Here we deploy these concepts in an analysis of 16S rRNA data to test the predictions in a wider range of environments.

## Results

We used data from the NCBI Reference Sequence (RefSeq) database separately and in combination with 16S-based community profiles. By processing these data (see Materials and Methods) we obtained *predicted proteomes*, which refer to proteomes for individual taxonomic groups in the RefSeq database that are based on automatic translation of genomic sequences, and *inferred community proteomes*, which represent the combination of predicted RefSeq proteomes with 16S-based microbial abundance profiles to estimate the amino acid compositions of proteomes for entire communities.

We analyzed the amino acid compositions of proteomes using two chemical metrics. Carbon oxidation state (*Z*_C_) of an organic molecule denotes the average charge on carbon atoms required for electroneutrality given formal charges of all other atoms. Also known by other names such as nominal oxidation state of carbon (NOSC), *Z*_C_ is an established metric for describing chemical transformations of natural organic matter (***Kroll et al., 2011***; ***Arndt et al., 2013***; ***Visconti and de Paz, 2021***), and has also been applied to analysis of metagenomic sequences (***Fones et al., 2019***; ***Lecoeuvre et al., 2021***). The equation for *Z*_C_ of proteins only requires elemental abundances (***Dick, 2014***; ***Dick et al., 2020***).

The second chemical metric, stoichiometric hydration state 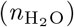, is computed from the number of water molecules in theoretical reactions to form proteins from a set of thermodynamic components. These components are also referred to as basis species, and provide a way of projecting elemental composition into chemical species such as O_2_ and H_2_O. Other than the requirement that they are the minimum number of species that can be linearly combined to represent all possible elemental compositions of the primary sequence of proteins, there is no thermodynamic constraint on the choice of basis species. A particular set of basis species – glutamine, glutamic acid, cysteine, O_2_, and H_2_O (abbreviated QEC) – was chosen to maximize the covariation between the independently computed *Z*_C_ of proteins and the number of O_2_ in their theoretical formation reactions (which are both measures of oxidation state) and to reduce that between *Z*_C_ and 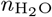 (***Dick et al., 2020***). These basis species have been used to identify distinct signatures of oxidation-reduction and hydration-dehydration reactions in both metagenomic sequences and clinical proteomic data (***Dick et al., 2020***; ***Dick, 2021b***).

To provide a taxonomic context for the analysis of 16S gene sequence datasets, the presentation of results begins with a visualization of the chemical metrics of the predicted proteomes of taxonomic groups in the RefSeq database.

### Chemical differences among taxonomic groups

Chemical metrics of predicted proteomes for phyla with the greatest number of representative lower-level taxa in the RefSeq database are plotted in ***Figure 1***. The first panel includes viruses and archaeal and bacterial phyla, and reveals that proteins in many viruses have a lower stoichiometric hydration state than most cellular organisms except for Bacteroidetes. The next panel excludes viruses to consider cellular organisms. The proteomes of organisms affiliated to Bacteroidetes have the lowest overall 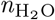, and those for Crenarchaeota, Fusobacteria, Actinobacteria, and Euryarchaeota, except for the classes Haloarchaea and Nanohaloarchaea, have relatively high 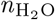. In terms of *Z*_C_, Actinobacteria, Planctomycetes, and Haloarchaea and Nanohaloarchaea within the Euryarchaeota are the groups with the most oxidized proteomes, whereas Crenarchaeota, Thermotogae, Fusobacteria, and Tenericutes have the most reduced proteomes. In the third panel, the stoichiometric hydration state distinguishes the main classes of Proteobacteria, with most orders of Alphaproteobacteria and Gammaproteobacteria at high and low 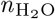, respectively, although some orders of the Alphaproteobacteria are considerably less hydrated. Notably, the proteobacterial class with the most reduced proteins is Epsilonproteobacteria. Members of this class are often identified in hydrothermal vent communities (***Nakagawa et al., 2005***; ***Campbell et al., 2006***), and in a proposed reclassification this class has been reassigned as a new phylum named Campylobacterota (***Waite et al., 2018***).

**Figure 1.**
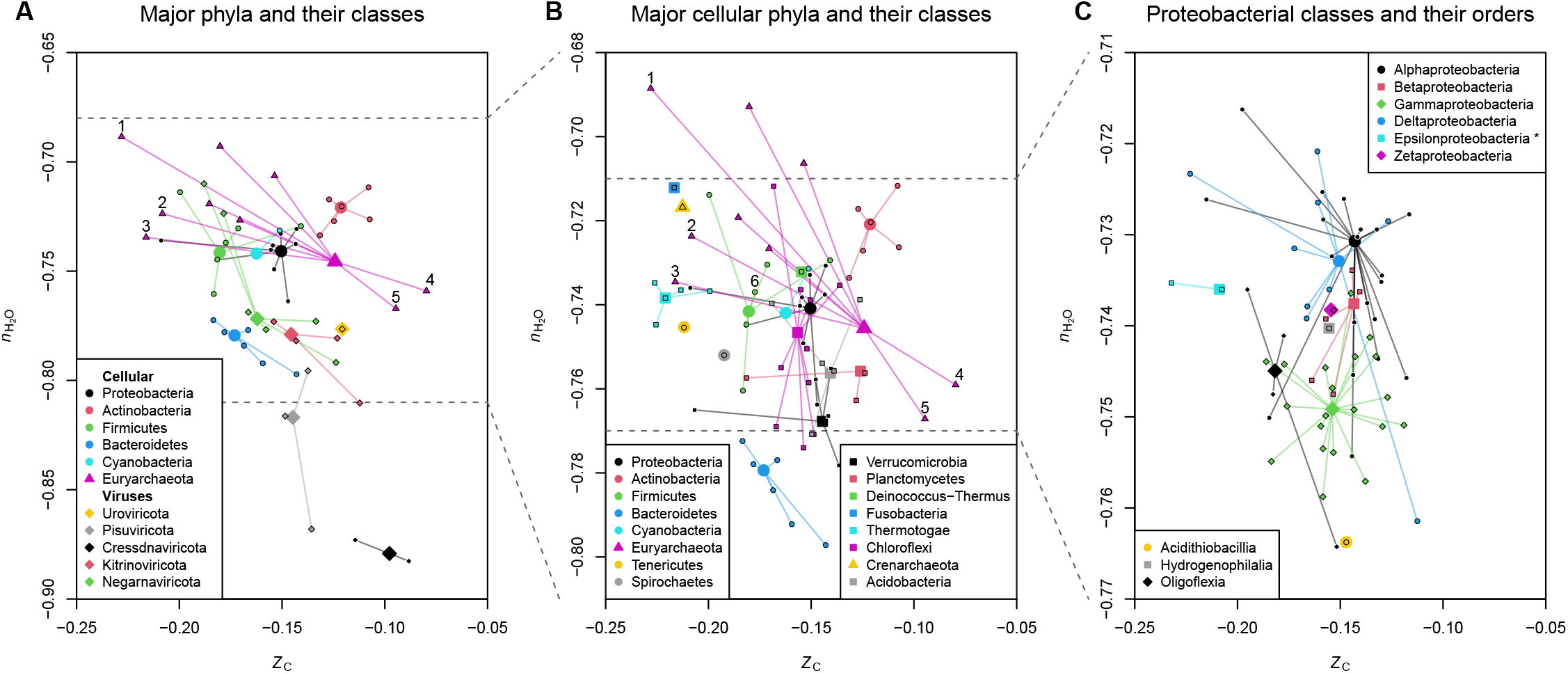
Distinct chemical parameters of predicted proteomes for major taxonomic groups. **(A)** Archaeal, bacterial, and viral phyla with more than 500 members at all lower levels (identified by unique taxonomic IDs in the NCBI taxonomy database); **(B)** archaeal and bacterial phyla with more than 60 lower-level members; **(C)** proteobacterial classes. Large symbols are for high-level taxa (phyla in **A** and **B**; classes in **C**) and small outlined symbols represent lower-level taxa (classes in **A** and **B**; orders in **C**). Points labeled 1, 2, 3, 4, and 5 are for the euryarchaeotal classes Methanococci, Archaeoglobi, Thermococci, Halobacteria, and Nanohaloarchaea, respectively, and the point labeled 6 is for the class Clostridia in the phylum Firmicutes. The taxonomic names are taken from the current NCBI taxonomy, but the Epsilonproteobacteria have recently been reclassified as Campylobacterota (phyl. nov.) (***Waite et al., 2018***). **Figure 1–source data 1**. Computed mean values of *Z*_C_ and 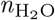 and names of all taxonomic groups shown in the plot, and numbers of individual taxa used to compute the means for each group.

The chemical representation of microbial proteomes generates many hypotheses about the effects of oxidation-reduction and hydration-dehydration reactions on biochemical evolution. For instance, the highly reducing environmental conditions of sediments and hydrothermal fluids (***Lovley and Klug, 1986***; ***Lin et al., 2016***) may have shaped the evolutionary processes that resulted in the low *Z*_C_ of proteins in the Methanococci, Archaeoglobi, and Thermococci, which are the most reduced classes in the Euryarchaeota and among the most reduced of all microbial classes shown in ***Figure 1***. In an even broader evolutionary context, the lower hydration state of viral proteomes than those of cellular organisms may not be surprising given the absence of a cytoplasmic compartment in viruses; in particular, the water content per gram of dry matter of geometrical viruses (those without a viral envelope) is considerably lower than that of *E. coli* cells (***Matthews, 1975***). The low 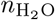 of Bacteroidetes, which are often enriched in suspended particles in seawater (***Fernández-Gómez et al., 2013***), aligns with our previous finding of lower 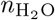 of proteins coded by metagenomes of particle-associated compared to free-living communities (***Dick et al., 2020***).

Some natural clustering is evident in ***Figure 1***, which indicates that the chemical composition of proteomes of classes within a phylum tend to be more similar to each other than to classes in other phyla. An interesting exception is the much higher *Z*_C_ and lower 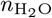 of Haloarchaea and Nanohaloarchaea than for other euryarchaeotal classes. The lower water activity associated with hypersaline environments may be one reason for these groups to have evolved proteomes with lower 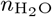. However, the high *Z*_C_ of these groups is most likely not an adaptation to more oxidizing conditions; indeed, the solubility of O_2_ decreases at higher salinity (***Sherwood et al., 1991***). Instead, the proteomes of many halophiles have greater numbers of acidic residues that stabilize the three-dimensional structure of proteins in high-salt conditions; because of the oxygen atoms contained in carboxylic acid groups, this adaptation also results in higher average oxidation state of carbon of the proteins (***Dick et al., 2020***).

### Microbial communities encode redox and salinity gradients

Published datasets from studies of hydrothermal systems, stratified water bodies, and microbial mats were selected to represent both local (within datasets) and global (across datasets) redox and salinity gradients. Accession numbers and literature references for the datasets analyzed here are listed in ***Table 1***.

**Table 1.**
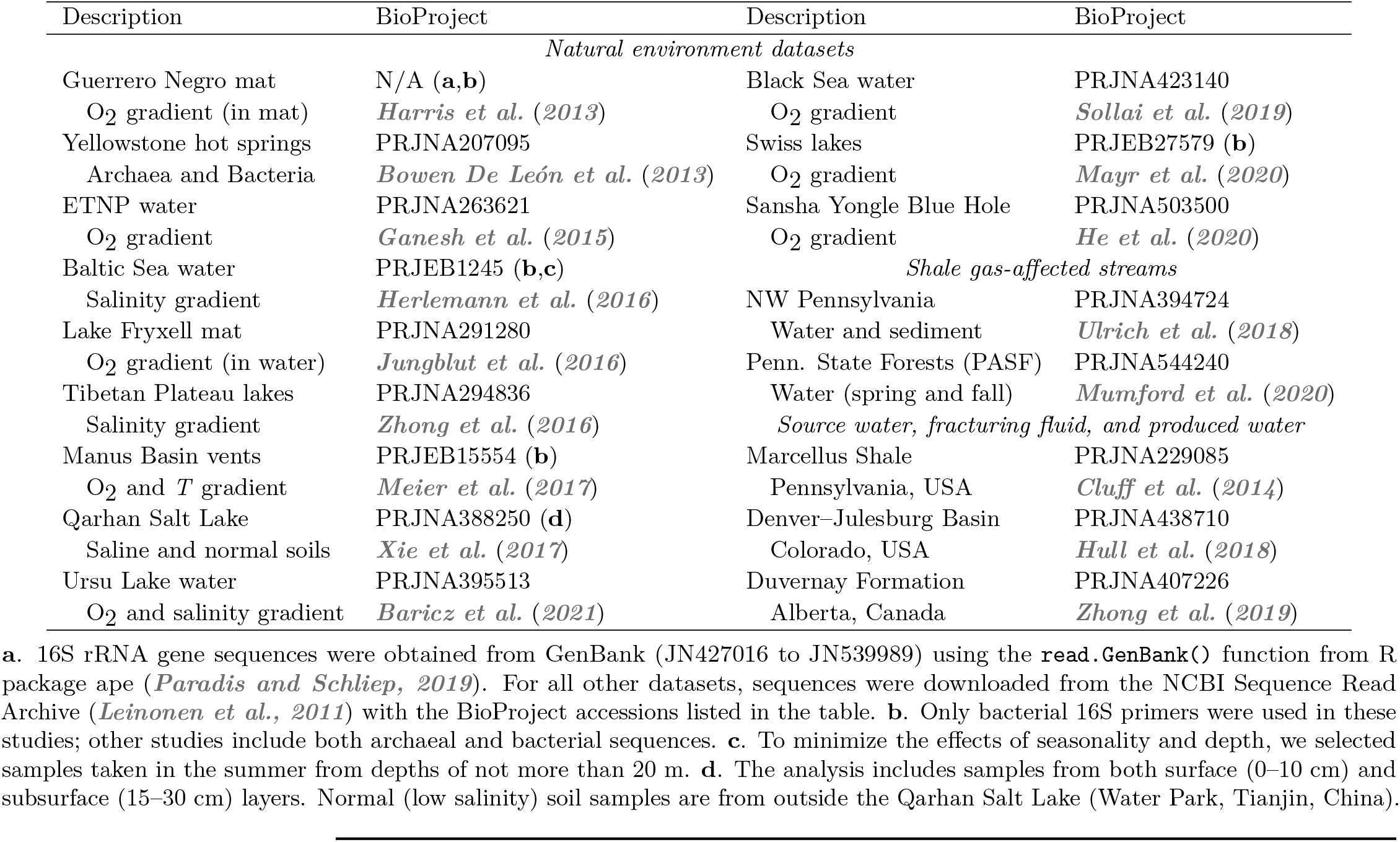
Sources of data used in this study.

| Description | BioProject | Description | BioProject |
| --- | --- | --- | --- |
| *Natural environment datasets* |  |  |  |
| Guerrero Negro mat | N/A (a,b) | Black Sea water | PRJNA423140 |
| O<sub>2</sub> gradient (in mat) | *Harris et al. (2013)* | O<sub>2</sub> gradient | *Sollai et al. (2019)* |
| Yellowstone hot springs | PRJNA207095 | Swiss lakes | PRJEB27579 (b) |
| Archaea and Bacteria | *Bowen De León et al. (2013)* | O<sub>2</sub> gradient | *Mayr et al. (2020)* |
| ETNP water | PRJNA263621 | Sansha Yongle Blue Hole | PRJNA503500 |
| O<sub>2</sub> gradient | *Ganesh et al. (2015)* | O<sub>2</sub> gradient | *He et al. (2020)* |
| Baltic Sea water | PRJEB1245 (b,c) | *Shale gas-affected streams* |  |
| Salinity gradient | *Herlemann et al. (2016)* | NW Pennsylvania | PRJNA394724 |
| Lake Fryxell mat | PRJNA291280 | Water and sediment | *Ulrich et al. (2018)* |
| O<sub>2</sub> gradient (in water) | *Jungblut et al. (2016)* | Penn. State Forests (PASF) | PRJNA544240 |
| Tibetan Plateau lakes | PRJNA294836 | Water (spring and fall) | *Mumford et al. (2020)* |
| Salinity gradient | *Zhong et al. (2016)* | *Source water, fracturing fluid, and produced water* |  |
| Manus Basin vents | PRJEB15554 (b) | Marcellus Shale | PRJNA229085 |
| O<sub>2</sub> and *T* gradient | *Meier et al. (2017)* | Pennsylvania, USA | *Cluff et al. (2014)* |
| Qarhan Salt Lake | PRJNA388250 (d) | Denver–Julesburg Basin | PRJNA438710 |
| Saline and normal soils | *Xie et al. (2017)* | Colorado, USA | *Hull et al. (2018)* |
| Ursu Lake water | PRJNA395513 | Duvernay Formation | PRJNA407226 |
| O<sub>2</sub> and salinity gradient | *Baricz et al. (2021)* | Alberta, Canada | *Zhong et al. (2019)* |

a. 16S rRNA gene sequences were obtained from GenBank (JN427016 to JN539989) using the `read.GenBank()` function from R package ape (*Paradis and Schliep, 2019*). For all other datasets, sequences were downloaded from the NCBI Sequence Read Archive (*Leinonen et al., 2011*) with the BioProject accessions listed in the table. b. Only bacterial 16S primers were used in these studies; other studies include both archaeal and bacterial sequences. c. To minimize the effects of seasonality and depth, we selected samples taken in the summer from depths of not more than 20 m. d. The analysis includes samples from both surface (0–10 cm) and subsurface (15–30 cm) layers. Normal (low salinity) soil samples are from outside the Qarhan Salt Lake (Water Park, Tianjin, China).

As visualized in ***Figure 2***, local redox gradients represented by stratified water bodies including the Black Sea (***Sollai et al., 2019***) and Lake Fryxell in Antarctica (***Jungblut et al., 2016***), submarine vents in the Manus Basin (***Meier et al., 2017***), and within the Guerrero Negro microbial mat (***Harris et al., 2013***) all provide evidence that carbon oxidation state is aligned with environmental redox conditions. *Z*_C_ is locally lower in the deep euxinic water of the Black Sea and anoxic water of Lake Fryxell, lower in the hotter water samples for the Manus Basin (which have greater input of reduced hydrothermal fluids), and lower just below the surface of the hypersaline Guerrero Negro microbial mat, which experiences a sharp oxygen gradient during the day (***Harris et al., 2013***). The Lake Fryxell dataset is also for mat samples, but the environmental gradient here is between relatively oxygenated shallow water and anoxic deeper water (***Jungblut et al., 2016***).

**Figure 2.**
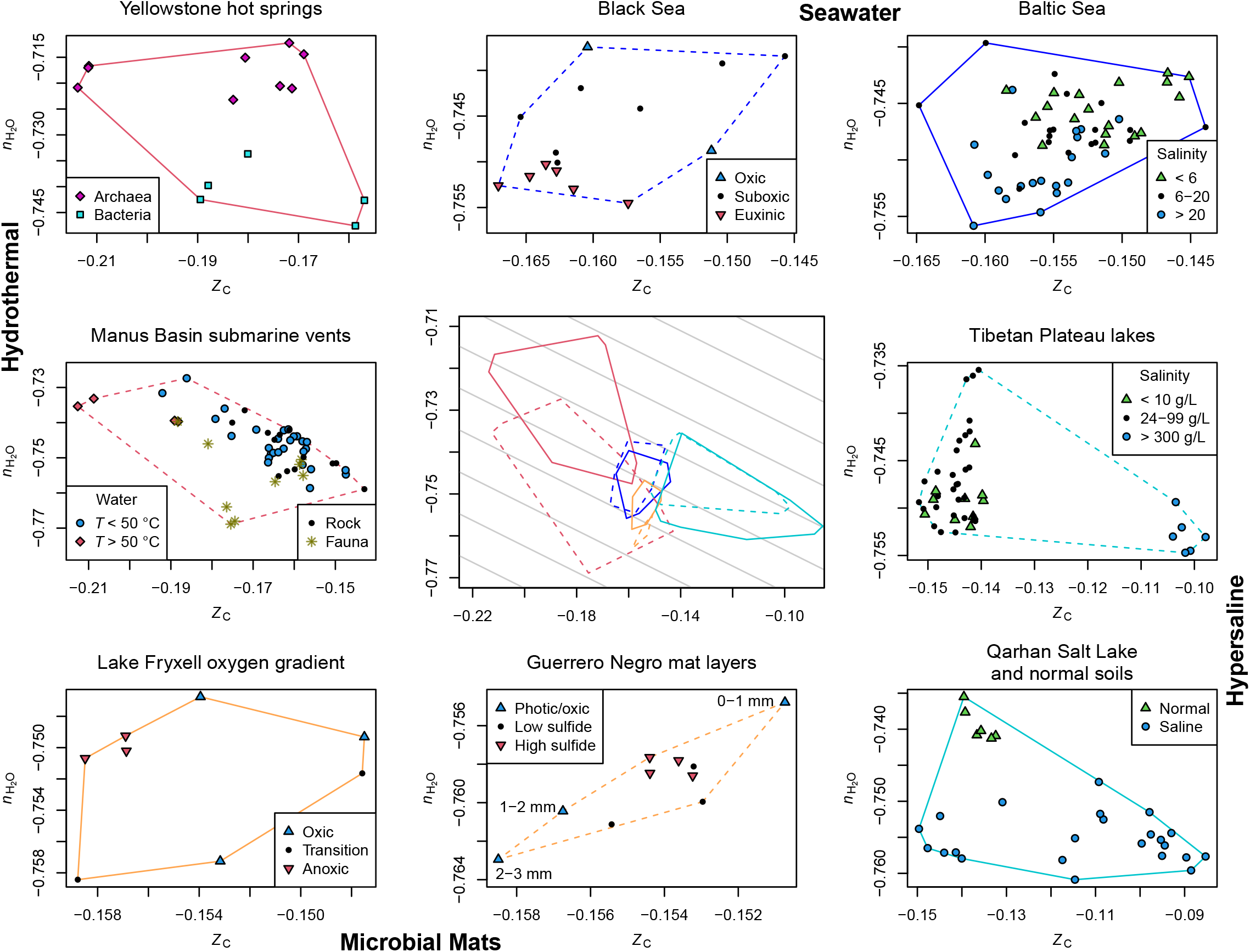
Inferred community proteomes from different environments have distinct chemical signatures. 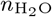 and *Z*_C_ were calculated for inferred community proteomes using 16S rRNA gene sequencing datasets for hydrothermal systems, seawater, hypersaline environments, and microbial mats. Sources of data are listed in ***Table 1***. The plot for each dataset shows individual samples as points and the convex hull containing all the samples. The convex hulls for individual datasets are assembled in the center index plot. The gray lines here have a slope corresponding to that of the regression between 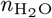 and *Z*_C_ for amino acids, and therefore represent the background covariation between these metrics when 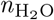 is calculated using the QEC basis species (glutamine, glutamic acid, cysteine, O_2_, and H_2_O) (***Dick et al., 2020***). The practical salinity values reported for the Baltic Sea (supplementary information of ***Herlemann et al., 2016***) have no units. **Figure 2–source data 1**. Computed values of *Z*_C_ and 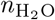, sample names, and SRA Run accessions used to make the plots.

Inferred proteomes of archaeal sequences from alkaline hot spring communities in Yellowstone National Park (***Bowen De León et al., 2013***) have higher 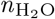 compared to bacterial sequences. This is aligned with expected phylogenetic differences, since the proteomes of major archaeal groups detected in these samples, including Crenarchaeota and Euryarchaeota (***Bowen De León et al., 2013***), generally have higher 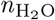 than bacterial proteomes (***Figure 1B***). Similarly, fauna samples in the Manus Basin dataset have lower 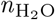 than the majority of fluid and rock samples, but in this case no archaeal sequences were reported, so differences in bacterial abundances are the reason for the observed chemical differences.

The Yellowstone and Manus Basin datasets represent hydrothermal systems that emit highly reduced fluids. In the index plot in the center of ***Figure 2***, it can be seen that these datasets are distributed toward lower *Z*_C_ than for the lake and seawater environments, thereby demonstrating that *Z*_C_ provides a signature of oxidation-reduction conditions on a global scale.

Both the Baltic Sea (***Herlemann et al., 2016***) and saline soils in the Qarhan salt lake (***Xie et al., 2017***) exhibit decreasing 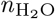 with greater salinity; we previously found a similar trend for proteins inferred from shotgun metagenomic sequences for various marine and freshwater environments (***Dick et al., 2020***). However, the trends for the Qarhan salt lake soils and Tibetan Plateau lake datasets (***Zhong et al., 2016***) are more strongly dominated by increasing *Z*_C_ in hypersaline conditions.

Many water bodies around the world develop vertical redox gradients as a result of microbial respiration of organic matter derived from the photic zone that leads to oxygen depletion with depth. Besides the Black Sea, we analyzed data for permanently stratified lakes in Switzerland (***Mayr et al., 2020***), the oxygen minimum zone of the Eastern Tropical North Pacific (ETNP) (***Ganesh et al., 2015***), the Sansha Yongle Blue Hole in the South China Sea (***He et al., 2020***), and Ursu Lake in Central Romania (***Baricz et al., 2021***). At each location, the *Z*_C_ of the inferred community proteomes generally decreases with depth (***Figure 3***). At the ETNP, the *Z*_C_ decreases strongly with depth in the free-living communities (0.2–1.6 µm size fraction), but to a lesser extent in particle-associated communities (1.6–30 µm size fraction). This might reflect environmental microniches and cell-cell interactions that to some extent reduce the sensitivity of these communities to external redox conditions.

**Figure 3.**
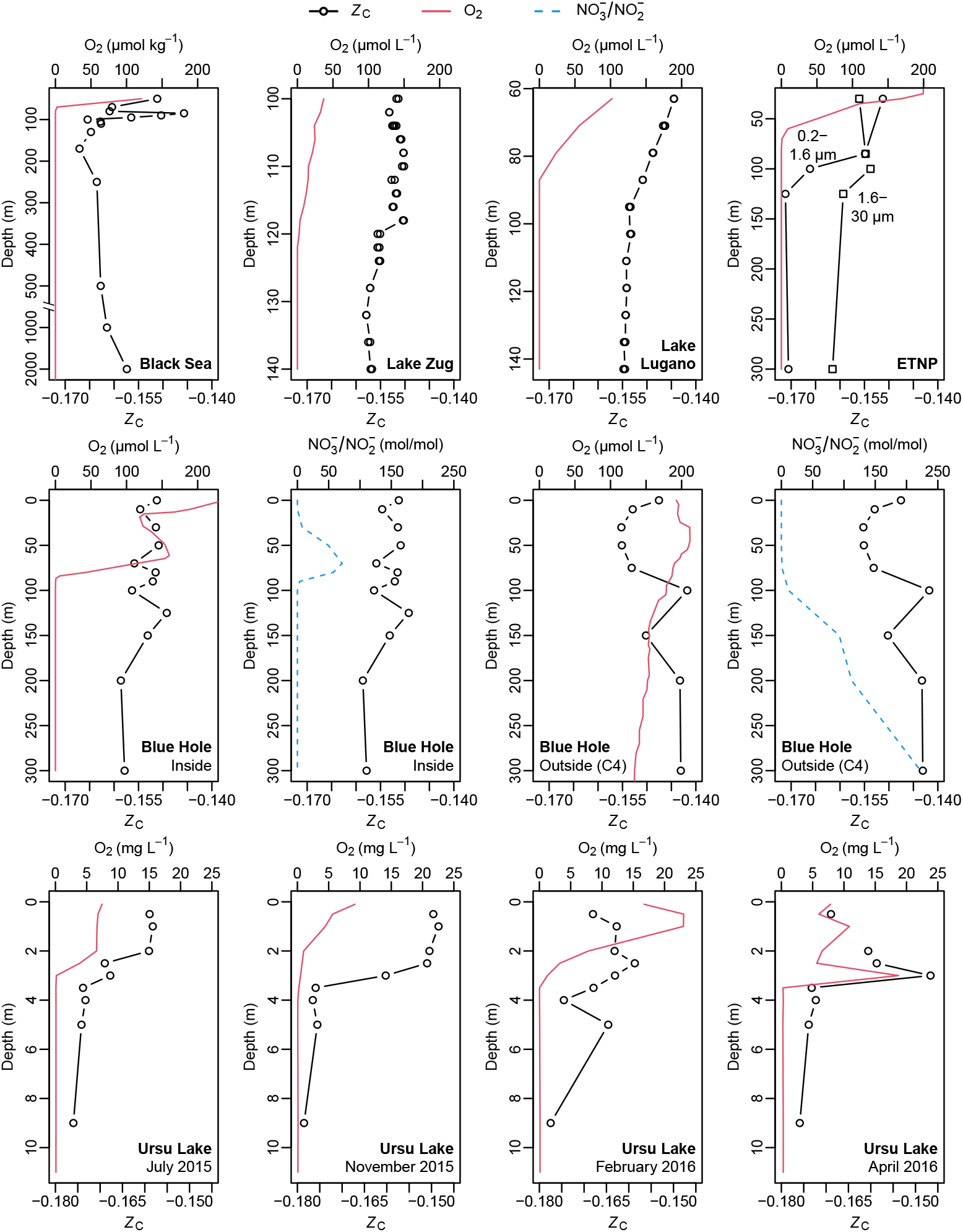
Lower carbon oxidation state is tied to oxygen depletion in water columns. All sites except for ETNP and station C4 outside the Blue Hole are permanently stratified. *Z*_C_ values were calculated for inferred community proteomes using published microbial 16S rRNA gene sequences (see **Table 1**). Oxygen concentrations were taken from the same publications, except for locations inside and outside the Blue Hole (***Xie et al., 2019***). For the Blue Hole, ratios of nitrate to nitrite (NO_3_^-^ / NO_2_^-^) are also plotted based on NO_3_^-^ and NO_2_^-^concentrations reported by ***Xie et al***. (***2019***). No *Z*_C_ value is shown for 1 m depth in Ursu Lake in April 2016 because fewer than 200 sequences remained for this sample after all sequence processing steps. **Figure 3–source data 1**. Computed values of *Z*_C_, chemical measurements taken from the literature, sample names and depths, and SRA Run accessions used to make the plots.

Although *Z*_C_ trends are dominated by oxygen gradients, a closer look suggests that other elemental redox systems may also contribute to the observed trends. There is a moderate rise in *Z*_C_ in the deepest (euxinic) layers of the Black Sea. Likewise, the low-to high-sulfide layers of the Guerrero Negro mat (below 3 mm) (***Harris et al., 2013***) have higher *Z*_C_ than the minimum at 3 mm (***Figure 2***). Therefore, despite the absence of oxygen, *Z*_C_ appears to rise locally in some sulfide-rich environments. At a non-stratified location outside the Sansha Yongle Blue Hole, the water remains oxygenated at depth. In the upper 100 meters, the profile of *Z*_C_ has a “C” shape, but maintains relatively high values at greater depths. In this case, the overall higher *Z*_C_ with depth appears to be more closely associated with the ratio of nitrate to nitrate than with O_2_ concentration.

Data from a study of Ursu Lake in Central Romania from summer 2015 to spring 2016 (***Baricz et al., 2021***) provide an opportunity to examine the temporal variability of oxygen concentration and chemical metrics of the microbial communities. At all times, both O_2_ and *Z*_C_ show an overall decline with depth. The profile of *Z*_C_ exhibits a broad maximum at around 2 m depth in November that becomes narrower and deeper through the winter and spring. The development of the *Z*_C_ maximum precedes that of an O_2_ maximum in February, and the two profiles exhibit a remarkable meter-scale correspondence in April.

### Physicochemical signal is strongest at high taxonomic levels

Not only the abundances but also the proteomic composition of individual taxonomic groups contribute to the overall chemical differences between microbial communities. To assess the chemical differences along redox and salinity gradients in more detail, the abundances and calculated *Z*_C_ or 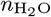 of major taxonomic groups at the domain, phylum, class, and genus levels are plotted in ***Figure 4*** for the Manus Basin and Baltic Sea. Because the lowest-level taxonomic classification for each sequence was used for mapping to the NCBI taxonomy to generate the inferred microbial proteomes (see Materials and Methods), within-group variation of *Z*_C_ and 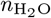 is possible above the genus level and is represented in ***Figure 4*** by sloping lines.

**Figure 4.**
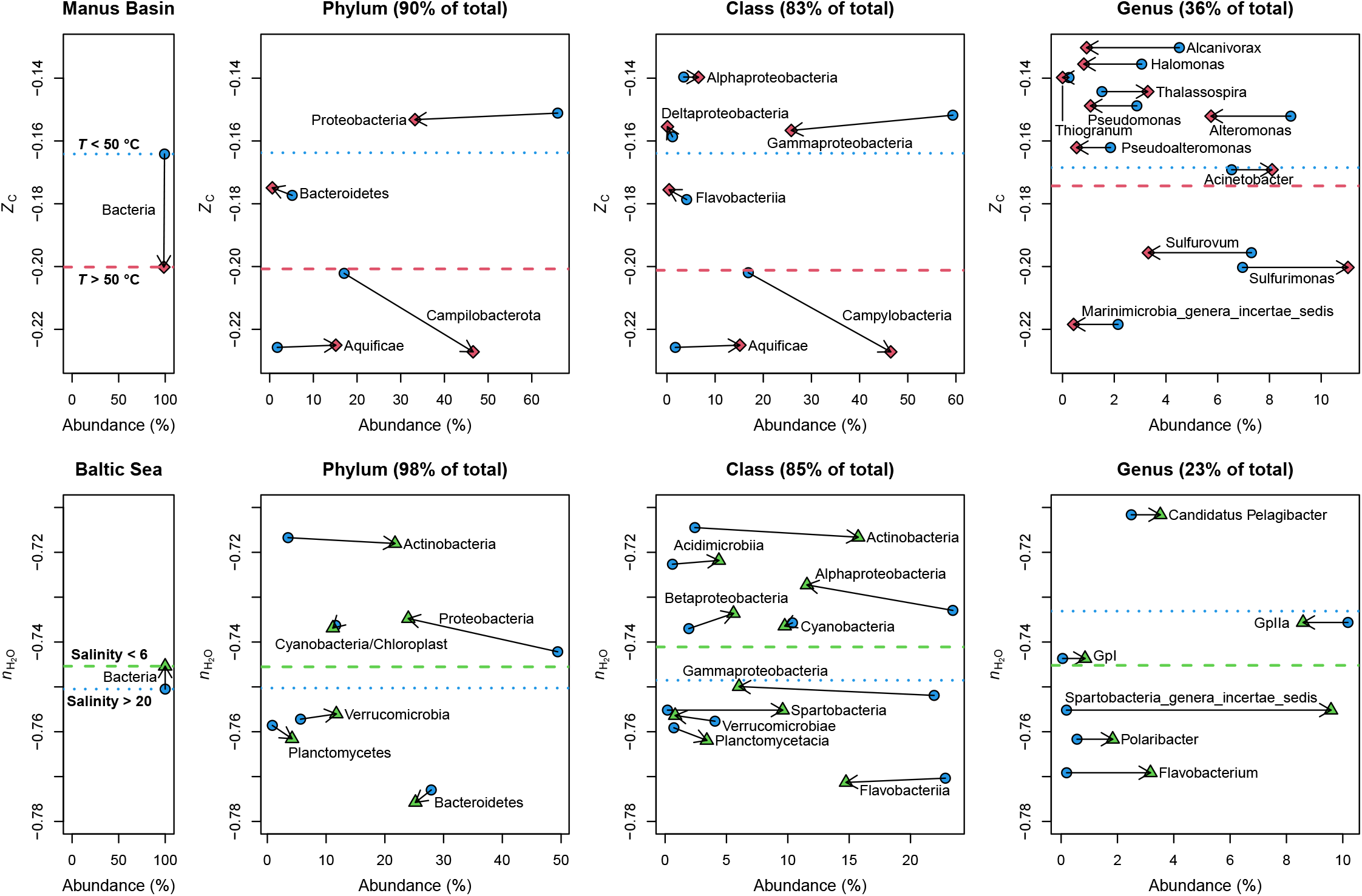
Changes of abundance and chemical composition at different taxonomic levels. Symbols represent taxon abundance-weighted mean values of *Z*_C_ for subsets of samples with relatively high and low temperature as a proxy for reducing and oxidizing conditions (*N* = 4 and 28, respectively) from the Manus Basin or 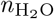 for subsets of samples with high and low salinity from the Baltic Sea (*N* = 19 and 18); environmental criteria used to allocate samples to subsets are shown in the plot. The leftmost plots represent all sequences classified by the RDP Classifier and mapped to the NCBI taxonomy at the domain level. In subsequent plots, sequences classified and mapped at lower taxonomic levels were combined to calculate the percentage abundance and *Z*_C_ or 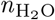 for each taxonomic group whose sequences make up at least 2% (for phylum or class) or 1% (for genus) of the total number of sequences from all samples. Percentages in the plot titles indicate the fraction of the whole community represented by groups shown in the plot. Arrows connect the same taxonomic group in different sample subsets and point to samples with higher temperature (Manus Basin) or lower salinity (Baltic Sea). Abundance-weighted means for all taxonomic groups shown in each plot are indicated by dashed lines for high temperature or low salinity samples and dotted lines for low temperature or high salinity samples. All taxonomic names are taken from the output of the RDP Classifier. Campilobacterota is probably a misspelling of the phylum name Campylobacterota (***Waite et al., 2018***).

At the phylum level, the high-temperature samples in the Manus Basin are associated with greater numbers of Aquificae and Campylobacterota (formerly Epsilonproteobacteria) and fewer Proteobacteria. These groups have relatively low and high *Z*_C_, respectively, which to a large extent explains the chemical difference at the whole-community level (leftmost plot). However, the campylobacterotal sequences themselves are affiliated with organisms whose proteomes have lower *Z*_C_ in the higher-temperature samples. Therefore, the whole-community chemical differences are due to both differential taxonomic abundances at the phylum level, which have the largest effect, as well as differential abundances within phyla. A similar finding applies to the major classes; at this level it is apparent that the proteobacterial contribution is mainly due to lower numbers of Gammaproteobacteria. The differential abundances of the identified genera yield a small *Z*_C_ difference between hotter and cooler fluids in the same direction as the whole-community trend.

Analogous reasoning can be used to interpret the trends in the Baltic Sea. The relatively high 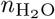 in low-salinity samples is mostly controlled by an increase in Actinobacteria. In contrast, Proteobacteria become less abundant at lower salinity, which to some extent counteracts the 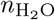 rise, but the within-group variation of Alphaproteobacteria, Betaproteobacteria, and Gammaproteobacteria is toward higher 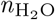at lower salinity. The genus-level assignments suggest an opposite trend (higher 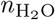 at higher salinity), but this is less likely to represent the actual differences because the low classification rate to the genus level (37%; see ***Table S1***) together with the 1% abundance cutoff for genera in ***Figure 4*** results in a low fraction of assignments represented at this level (23%).

### Application to datasets for shale gas operations

During hydrocarbon extraction from unconventional reservoirs (shale gas), hydraulic fracturing fluid is injected into shale formations to create extensive horizontal fracture networks that improve hydrocarbon recovery. The injected fluid mixes and reacts with natural formation waters and the fractured rock surfaces. Water that returns to the surface is initially referred to as flowback and later as produced water during the hydrocarbon production stage of the well (***Khan et al., 2021***). Chemical oxidants are commonly added to the injected fracturing fluid; these additives enhance mineral dissolution and can also have biocidal effects. Even without such additives, fracturing fluids generally consist of large amounts of water from surface sources, which makes them highly oxidized compared to the reducing conditions in organic-rich shale.

Changes in the chemistry and biology of streams that are in proximity to or may be affected by shale gas operations are an important issue for environmental assessments of shale gas operations. The chemistry of affected surface streams is thought to reflect the possible input of methane and/or chemical additives in fracking fluid from nearby wells, though the overall strength of these associations has been debated (***Mumford et al., 2020***).

Other authors have noted that little quantitative information is available about the changes in redox conditions of flowback and produced fluid over time (***Mouser et al., 2016***; ***Liden et al., 2019***). In one case, oxygen was found to be rapidly depleted in flowback and produced waters in the Duvernay Formation in Alberta, Canada (***Zhong et al., 2019***). In a study on the Marcellus Shale in Pennsylvania, USA, the abundances of S-bearing organic compounds determined from FT-ICR-MS measurements exhibited a decrease in carbon oxidation state compared to injected fluids (***Luek et al., 2019***). Furthermore, oxidation-reduction potential (ORP) measured using a multiprobe is lower (more reducing) in groundwater samples with higher concentrations of CH_4_ (***LeDoux et al., 2016***). Therefore, we predicted that inferred community proteomes in produced water would be shifted toward a more reduced state compared to source water. A related prediction is that putative hydrocarbon input to surface streams has a similar reducing effect, but the effect may be smaller because of the greater extent of dilution.

Analysis of 16S sequence data for water from streams in Northwestern Pennsylvania (***Ulrich et al., 2018***) shows that both *Z*_C_ and 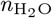 are lower in streams affected by Marcellus Shale activity compared to unaffected streams (***Figure 5A***). Smaller differences, but in the same direction, are found for stream sediments in the same study and for stream water in another study (***Mumford et al., 2020***) (***Figure 5B***). Much larger shifts, toward lower *Z*_C_ and higher 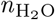, occur in flowback and produced waters compared to source waters and injected fracturing fluids. This trend is evident for not only the Marcellus Shale (***Cluff et al., 2014***) but also the Denver–Julesburg Basin in Colorado, USA (***Hull et al., 2018***) and the Duvernay Formation (***Zhong et al., 2019***) (***Figure 5C*** and ***Figure 5D***). Notably, the magnitudes of the *Z*_C_ differences in the Denver–Julesburg Basin and Duvernay Formation are the largest among all datasets considered in this study, and the Marcellus Shale has a larger negative *Z*_C_ compared to all datasets except for the Manus Basin hydrothermal vents (***Figure 6***). Therefore, highly reducing conditions associated with subsurface or hydrothermal systems are manifested as great differences in carbon oxidation state of inferred community proteomes, whereas oxygen gradients in water bodies exhibit a smaller *Z*_C_.

**Figure 5.**
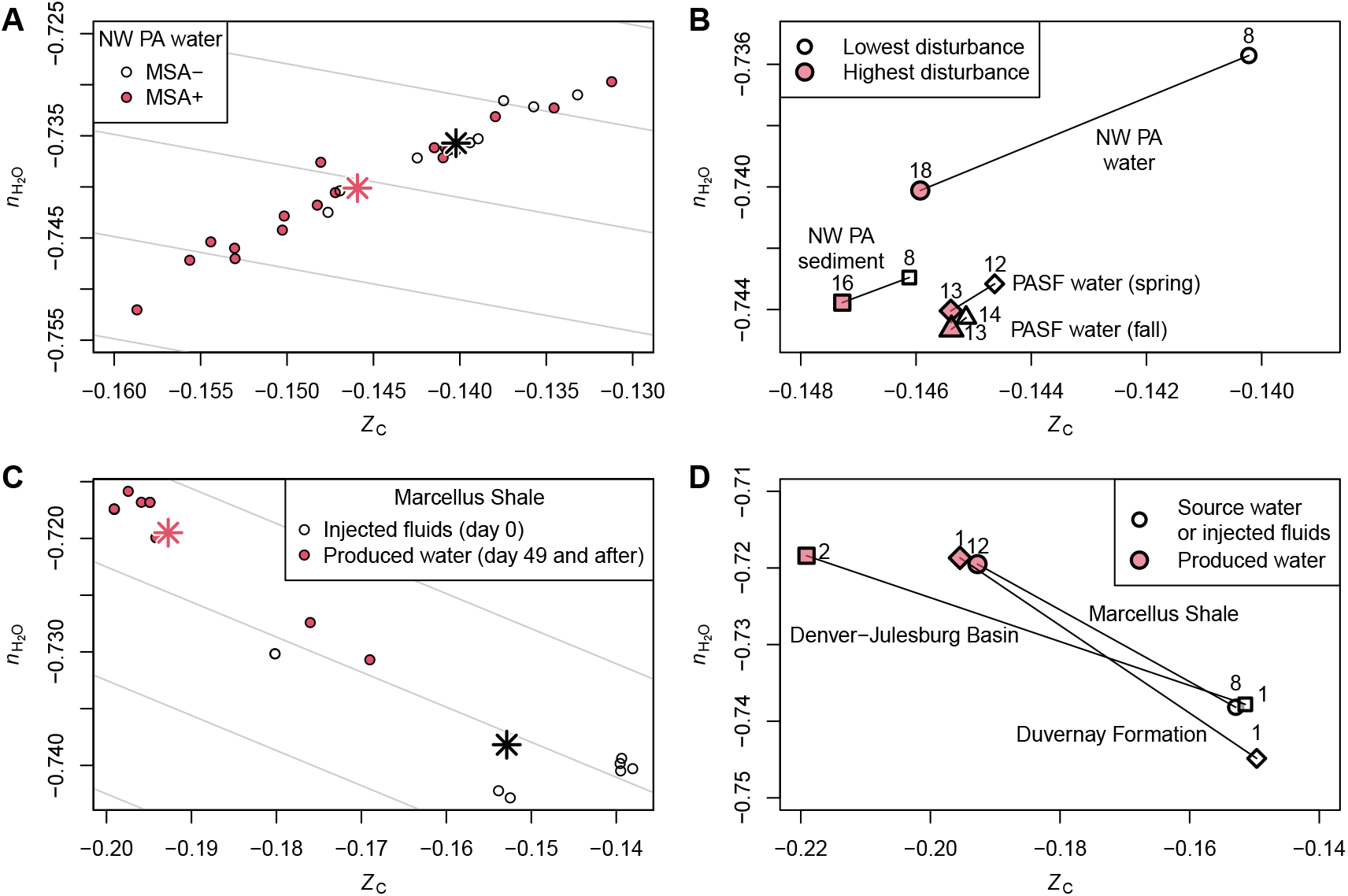
Decreased carbon oxidation state of inferred proteomes for communities affected by shale gas extraction. **(A)** Water samples from streams affected by Marcellus Shale activity (MSA+) and non-affected streams (MSA-) in Northwestern Pennsylvania (***Ulrich et al., 2018***); star-shaped symbols represent group means. **(B)** Mean values for sample groups in various studies of streams in Pennsylvania: Northwestern Pennsylvania (water and sediment samples) (***Ulrich et al., 2018***) and Pennsylvania State Forests (PASF; water samples in spring and fall) (***Mumford et al., 2020***). **(C)** Injected fluids and produced water from a hydraulically fractured well in the Marcellus Shale (***Cluff et al., 2014***); star-shaped symbols represent group means. **(D)** Mean values for samples of produced water compared to injected fluids or source water for hydraulically fractured wells in the Marcellus Shale (***Cluff et al., 2014***), Denver–Julesburg Basin (***Hull et al., 2018***), and Duvernay Formation (***Zhong et al., 2019***). Numbers of runs used for calculating group means are shown next to points in **(B)** and **(D)**; for the Marcellus Shale, there are two runs for each biological sample. In this figure, data for the samples collected in 2014 for NW PA water and sediment (***Ulrich et al., 2018***) are used; in ***Figure 6*** water samples from all available years (2012 to 2016) are used. **Figure 5–source data 1**. Computed values of *Z*_C_ and 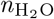, SRA Run accessions, and sample names and types (e.g. MSA+ or MSA-, or source or produced water) used for calculating group means.

**Figure 6.**
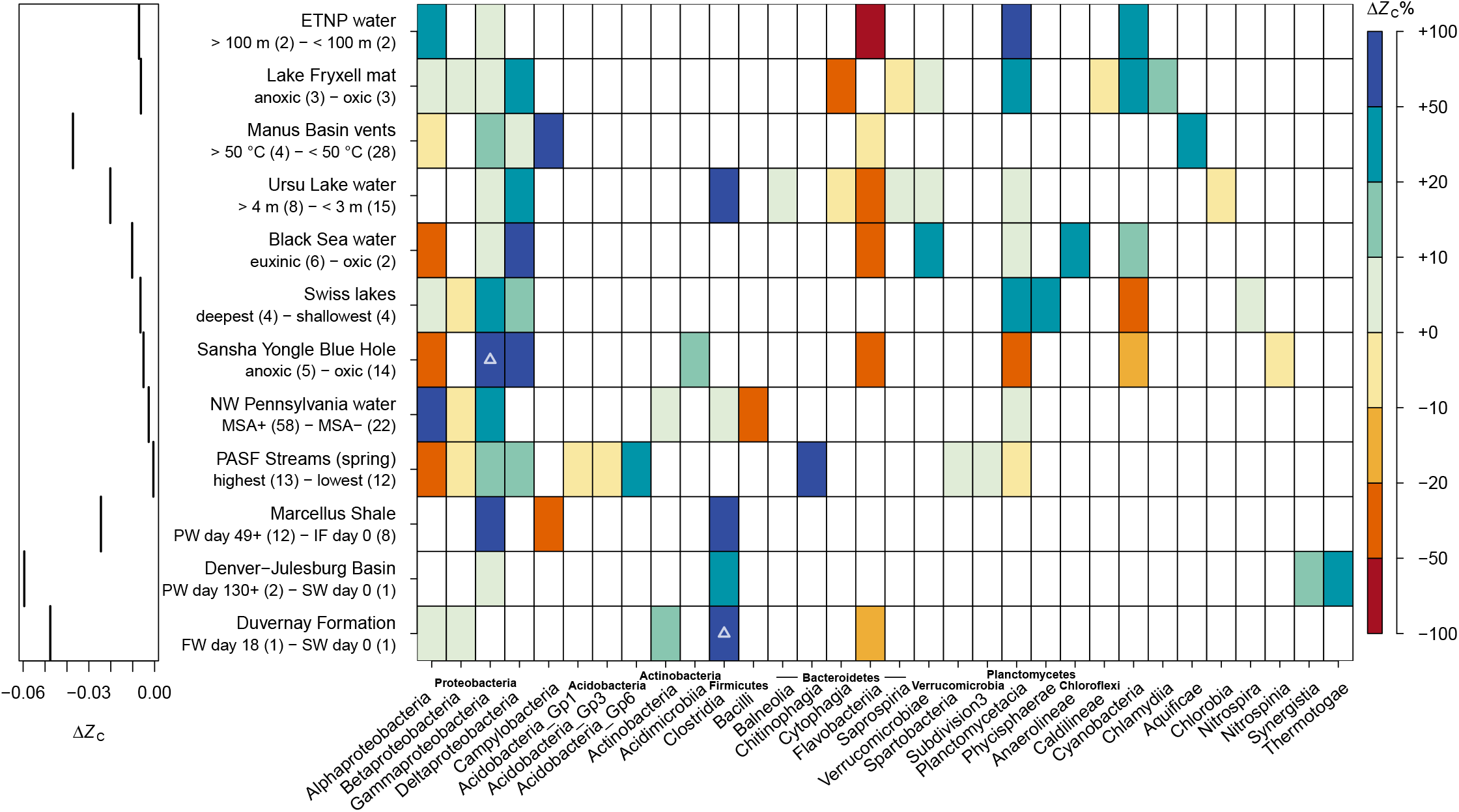
Contributions of taxonomic classes to the difference in carbon oxidation state of inferred community proteomes between relatively oxidized and reduced communities. Samples in each dataset were categorized into oxidized and reduced subsets as indicated in the dataset descriptions; numbers of samples in each category are given in parentheses. For each dataset, colored boxes indicate classes that are at least 2% abundant across all included samples. Inferred community proteomes for the oxidized and reduced subsets of samples were aggregated and used to calculate values of Δ*Z*_C_. The color scale represents the contribution of each class to the overall Δ*Z*_C_, calculated using ***Equation 1***. Positive percentages mean that the change in abundance and/or *Z*_C_ of the class itself is in the same direction as the overall Δ*Z*_C_ (i.e. more negative in reducing conditions). Negative percentages indicate contributions by classes that are in the opposite direction. The color scale is limited to percentages in the range [-100, 100]; values > 100 are indicated by triangles inside the boxes. The sum of percentages for each dataset is 100. Datasets representing redox gradients were selected from ***Table 1*** and are ordered as in the table. Datasets for NW Pennsylvania sediment and PASF Streams (fall) were omitted because they have a very small Δ*Z*_C_ at the class level. Classes are grouped by phylum but their ordering is otherwise arbitrary. Abbreviations: FW – flowback water; IF – injected fluids; PW – produced water; SW – source water. MSA+, MSA-, highest, and lowest are terms used by previous authors to describe the level of disturbance of streams from unconventional hydrocarbon extraction (***Ulrich et al., 2018***; ***Mumford et al., 2020***). Because of the different sample depths and O_2_ profiles of the Swiss lakes (***Figure 3***), the deepest and shallowest samples from both Lake Zug and Lake Lugano were selected. **Figure 6–Figure supplement 1**. Plots of abundance and *Z*_C_ for taxonomic classes in oxidized and reduced sample subsets.

## Discussion

The main finding of this study is that the oxidation state of inferred community proteomes decreases in more reducing conditions at global and local scales. Closer examination of selected datasets (***Figure 4***) indicates that the whole-community chemical differences are mainly associated with changes in abundances of particular phyla and that within-group variation of particular classes is often in the same direction, indicating that physicochemical conditions shape microbial communities at multiple taxonomic levels.

The results here are consistent with our earlier analysis of shotgun metagenomic data that showed lower *Z*_C_ of proteins in Yellowstone hot springs and the Lost City hydrothermal vents compared to ambient seawater (***Dick et al., 2019***). Similarly, ***Lecoeuvre et al. (2021***) analyzed shotgun metagenome sequences from the newly discovered Old City hydrothermal field. The mean value they reported for *Z*_C_ of proteins (−0.168) lies within the range for hydrothermal systems obtained here (***Figure 2***) and is lower than all but the most reduced community proteomes inferred for anoxic waters (***Figure 3***). Therefore, *Z*_C_ values calculated using inferred community proteomes (this study) or from shotgun metagenomic sequences (***Dick et al., 2019***; ***Lecoeuvre et al., 2021***) appear to be commensurate.

In our previous analysis of shotgun metagenomic data, we found conflicting trends for *Z*_C_ in oxygen minimum zones (***Dick et al., 2019***). We speculated that preferential degradation of low-GC content extracellular DNA by heterotrophs could leave behind resistant genes that are more likely to undergo horizontal gene transfer; the putative enrichment of AT-rich genes would be manifested by higher *Z*_C_ of the inferred proteins (***Dick et al., 2019***). It has also been proposed that horizontal gene transfer could hinder the use of shotgun sequences for accurate taxonomic identification (***Tessler et al., 2017***), although we note that our previous analysis of *Z*_C_ of shotgun sequences was not based on taxonomic assignments. In contrast to the results for shotgun metagenomes, the present study uncovered a strong signal of decreasing *Z*_C_ with depth in multiple 16S rRNA datasets for stratified water bodies.

The highly saline produced waters from many shales converge toward a common profile dominated by the halophilic and anaerobic *Halanaerobium* (***Mouser et al., 2016***), but in the Denver–Julesburg Basin *Thermoanaerobacter*, which has similar metabolic capabilities, is present instead (***Hull et al., 2018***). The predicted RefSeq proteomes of these groups have *Z*_C_ values of -0.199 and -0.227, respectively; the very low oxidation states of these abundant groups, which are both members of Clostridia (see ***Figure 1B***) accounts for much of the decrease in *Z*_C_ in produced fluids. Notably, hypersaline conditions in other environments including lakes and soils are characterized by relatively high *Z*_C_ (this study and ***Dick et al., 2020***), so the finding of an opposite trend for shale gas wells strengthens the interpretation that reducing conditions are a primary driver of community structure in produced water.

More work is needed to derive a chemical metric that captures the relationship between community composition and salinity. Because of osmotic forces, higher salinity should have a dehydrating effect, but this prediction is not supported by the increase of 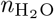 of inferred community proteomes that is observed for produced fluids. Higher 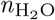 may be intrinsically linked to lower *Z*_C_ as a result of the background correlation between these metrics when 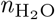 is calculated using the QEC basis species (gray lines in ***Figure 5A*** and ***Figure 5C***; see also ***Dick et al., 2020***). However, the relations between *Z*_C_ and 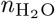 are not universally dictated by the basis species, as shown by the positive slopes between datasets for stream communities in ***Figure 5B***.

At this point it is not possible to predict with this method which taxa are actually present in a community. Nevertheless, analysis of the contribution of individual classes to the differences in *Z*_C_ within datasets reveals some provocative patterns for these taxa that are present (***Figure 6***). In all datasets where they make a major contribution to the oxidation-reduction differences, the Gammaproteobacteria, Deltaproteobacteria, and Clostridia contribute to lower the overall *Z*_C_ of the communities in more reducing environments, due to either changes in abundance, within-group *Z*_C_, or both (see ***Figure 6–Figure Supplement 1*** for a depiction of the values used for these calculations). These groups can therefore be regarded as the microbial drivers of the environmental shaping by redox gradients. In contrast, the change in abundance of Flavobacteriia opposes the community-level Δ*Z*_C_ for many datasets; a similar trend is also apparent for Cytophagia, which is another group within the Bacteroidetes, in some other datasets. These classes have relatively reduced proteomes (i.e. more negative *Z*_C_), but they tend to be more abundant in more oxidizing environments, which explains their opposing contribution (***Figure 6–Figure Supplement 1***). Therefore, these classes can be described as detracting from the shaping of various communities by local redox gradients. To what extent the phenotypic traits of Bacteroidetes, known for degradation of high molecular weight organic matter (***Fernández-Gómez et al., 2013***), may underlie the unique behavior of these groups remains unknown; whether these trends could be related to the very low 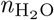 of their proteomes (***Figure 1***) is also an open question.

Our findings for shale gas systems suggest that in addition to salinity, the steep redox gradient between the surface (oxidizing) and subsurface (reducing) is a primary factor that shapes the structure of microbial communities. Other authors have commented on the dearth of redox and oxygen measurements in samples collected from black shale well sites (***Mouser et al., 2016***), and a lack of such measurements is also apparent in the USGS National Produced Waters Geochemical Database (***Blondes et al., 2018***). Of the 114943 records in the database, there are only 66 with measurements of dissolved oxygen (O_2_) or ORP, and these are only for conventional or geothermal wells. The multivariate analyses performed in previous studies (***Cluff et al., 2014***; ***Hull et al., 2018***; ***Zhong et al., 2019***) identified variables including dissolved inorganic and organic carbon, salinity, pH, and temperature as major environmental drivers of the differences in microbial community composition. Because oxygen or redox measurements of produced waters from unconventional wells were not available, those analyses could not have been used to identify an association between microbial communities and oxidation-reduction state of the fluids. However, such an association is exactly what is predicted by the chemical analysis in this study.

In spite of the oxygenation of injected fluids, subsurface conditions are likely to be anoxic (***Zhong et al., 2019***), so other types of measurements should be considered for monitoring the redox state of produced water. For instance, the USGS database cites a study from 2009 that gives both NO_3_^-^ and NO_2_^-^ measurements in injected and produced water from the Marcellus Shale (***Hayes, 2009***). Monitoring the dynamics of N species in conjunction with biological sampling would give further insight into the extent of nitrate reduction in the subsurface (***Orcutt et al., 2011***; ***Cluff et al., 2014***) and could also serve as a proxy for *in situ* redox conditions through a metric such as the NO_3_^-^/NO_2_^-^ ratio (***Ducluzeau et al., 2014***). ORP is another measure of redox conditions; however, they are generally considered more difficult to interpret because the electroactivity of chemical species in the sample is affected by kinetic barriers that affect the potential reading but are not readily quantified. Nevertheless, ORP measurements can yield information about electrochemical reactions that are relevant to microbial growth, especially if the measurements are made continuously in time (***Markelova et al., 2017***).

By leveraging the chemical information contained in genomic sequences, it is possible to achieve a broader view of the coupling between inorganic and organic oxidation-reduction reactions that is essential for all ecosystems (***Burgin et al., 2011***; ***Orcutt et al., 2011***). Analysis of inferred community proteomes indicates that redox gradients have the strongest effect on microbial communities in produced fluids, followed by hydrothermal environments and oxygen gradients in stratified water bodies. The results suggest that more comprehensive monitoring of dissolved oxygen concentrations or other redox indicators should be used to better characterize the responses of microbial communities in produced water and streams affected by unconventional hydrocarbon extraction, and ecosystems in general.

## Materials and Methods

### Data sources, processing, and classification

16S rRNA gene sequences were downloaded from the NCBI Sequence Read Archive (SRA) except for the Guerrero Negro microbial mat sequences (***Harris et al., 2013***), which were obtained from GenBank. The processing pipeline consisted of merging of paired-end reads, length and quality filtering, removal of singletons, subsampling, chimera removal, and taxonomic classification. VSEARCH version 2.15.0 (***Rognes et al., 2016***) was used to merge Illumina paired-end reads; for some datasets with low-quality reverse reads, only the forward reads were used as in previous studies (***Ulrich et al., 2018***). For Illumina datasets, quality and length filtering were done with the options `-fastq_maxee_rate` 0.005 (i.e. maximum one expected error for every 200 bases) and `-fastq_trunclen` with a length value depending on the specific dataset. For 454 datasets, where reads are generally longer but have more variable length, quality and length filtering were done with `-fastq_truncqual 15 -fastq_minlen 200 -fastq_maxlen 600;` also, the option `-fastq_stripleft 18` was used to remove adapter sequences. Sequence processing statistics and additional details are given in ***Table S1***.

After filtering, the remaining sequences for all samples in each dataset were pooled and singletons (sequences that appear exactly once, but not including subsequences of other sequences) were removed. Then, samples were subsampled to a depth of 10000 sequences; samples with fewer than 10000 sequences were not affected. The subsampling reduces the processing time for chimera detection, which is the longest step in the pipeline, and retains enough sequences for classifying the major taxonomic groups in the communities. Reference-based chimera detection was performed using the VSEARCH command `-uchime_ref` with the SILVA 138.1 SSURef NR99 database (***Quast et al., 2012***). Sequences identified as chimeras or borderline chimeras were removed.

The remaining sequences were processed with the Ribosomal Database Project (RDP) (***Cole et al., 2014***) Classifier version 2.13 (***Wang et al., 2007***) with the provided training set (RDP 16S rRNA training set No. 18 07/2020). The sequence classifications for all samples in each dataset were merged using the RDP Classifier command `merge-count`.

### RefSeq proteomes

Predicted protein sequences were obtained from the RefSeq database release 206 (2021-05-21) (***O’Leary et al., 2016***) for all 49448 bacterial, archaeal and viral taxa identified by a unique taxonomic ID (taxid). For each taxid, the available taxonomic names at ranks of superkingdom, phylum, class, order, family, genus, and species were parsed from the current (as of the RefSeq release date) NCBI taxonomy files, and the amino acid compositions of all available protein sequences were summed and divided by the number of proteins to generate the average amino acid composition of the predicted proteome. To calculate the amino acid compositions for taxonomic groups at genus and higher levels, those for each taxon within this group, including lower levels, were summed then divided by the number of taxa. The number of proteomes, corresponding to the number of taxonomic groups at each level, is 4788 (genus), 763 (family), 303 (order), 140 (class), 78 (phylum), and 3 (superkingdom).

### Taxonomy mapping

To infer the amino acid compositions of communities from 16S sequencing data, RDP classifications at only the root or domain level were first omitted because they provide very little taxonomic resolution. Sequences assigned to RDP class- and family-level name Chloroplast or genus-level names Chlorophyta and Bacillariophyta were also discarded because they do not fall within the archaeal and bacterial taxonomy used by NCBI. All remaining classifications were retained for attempted mapping to the NCBI taxonomy. The lowest-level classification returned by the RDP classifier for each sequence was used; the classification rate to genus level varies widely among datasets (***Table S1***) so higher-level classifications were used where needed.

In general, mapping from the RDP Classifier to the NCBI taxonomy was performed by text matching of both the taxonomic rank and name. Some particular mappings were used to improve the representation of common taxa in the datasets. The RDP phylum Cyanobacteria/Chloroplast, class Planctomycetacia, and genus *Escherichia/Shigella* were mapped to the NCBI phylum Cyanobacteria, class Planctomycetia, and genus *Escherichia*, respectively. The RDP order Rhizobiales was mapped to the NCBI order Hyphomicrobiales (***Hördt et al., 2020***). The RDP taxon Spartobacteria genera *incertae sedis*, which is relatively abundant in the Baltic Sea (***Herlemann et al., 2016***), was mapped to NCBI class Spartobacteria. The RDP taxon Subdivision3 genera *incertae sedis*, which was identified here in some stream datasets (***Mumford et al., 2020***) was mapped to NCBI family Verrucomicrobia subdivision 3. The RDP taxon Marinimicrobia genera *incertae sedis*, which was identified in this study in some deep ocean datasets (***Ganesh et al., 2015***; ***Meier et al., 2017***), was mapped to NCBI species Candidatus Marinimicrobia bacterium, which is the only representative of the Candidatus Marinimicrobia phylum in the RefSeq database. Among Acidobacteria, which are fairly abundant in river water and sediment, the RDP genus-level classifications Gp1 and Gp6 were mapped to NCBI genera *Acidobacterium* and *Luteitalea*, respectively, which are members of Acidobacteria subdivisions 1 and 6 (***Eichorst et al., 2018***). The RDP genus-level cyanobacterial groups GpI, GpIIa, and GpVI were mapped to the NCBI genera *Nostoc, Synechococcus*, and *Pseudanabaena*, and the RDP taxon Family II was mapped to the family Synechococcaceae; these mappings are based on names of members of these groups given in Bergey’s Manual (***Castenholz et al., 2001***), although the mappings are necessarily imperfect because of inconsistencies with the NCBI taxonomy.

Any other RDP classifications whose rank and name could not be matched to the NCBI taxonomy were removed from the subsequent calculations. Note that sequences were mapped using the lowest available taxonomic level in the output of the RDP Classifier; this is why the phylum Campilobacterota (the name output by the RDP Classifier) is represented in ***Figure 4*** although no explicit mapping to the NCBI name Campylobacterota was done. Across all datasets, a median of 96.2% of RDP classifications at all levels from genus to phylum were mapped to the NCBI taxonomy (***Table S1***). The lowest classification rate is for a dataset for produced water from shale gas wells (***Hull et al., 2018***), in which the genus *Cavicella* makes up 27% of the RDP classifications but has no counterpart in the available RefSeq proteomes (***Table S2***).

### Analysis of chemical composition

Values of *n*_C_, *Z*_C_ and 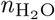 were computed for each taxonomic group in RefSeq whose amino acid compositions were obtained as described above, using the `ZCAA()` and `H2OAA()` functions in the canprot package (***Dick, 2021b***) and a modified function to calculate *n*_C_ (number of carbon atoms). For individual or aggregated samples, the RDP counts for each mapped taxon were multiplied by 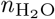 of that taxon, summed, and divided by the total counts to obtain the abundance-weighted mean 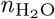 for the inferred community proteome. For *Z*_C_, the mean was additionally weighted by carbon number as described previously (***Dick et al., 2020***): *Z*_C_ = Σ*n*_*i*_*n*_C,*i*_*Z*_C_*/*Σ*n*_*i*_*n*_C,*i*_, where *n*_*i*_, *n*_C,*i*_ and *Z*_C,*i*_ are the count, carbon number, and carbon oxidation state of the *i*th taxon. Because *Z*_C_ is a per-carbon quantity and 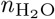 values are per-residue, comparisons can be made between proteomes with different average protein length.

To compare subsets of samples within datasets, RDP classifications for all samples in each subset were first aggregated, then *Z*_C_ and 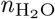 were calculated from the aggregated taxon counts. The percent contribution of the *i*th taxon (Δ*Z*_C_%) to the difference in *Z*_C_ between sample subsets “o” and “r” (for oxidized and reduced) was calculated from

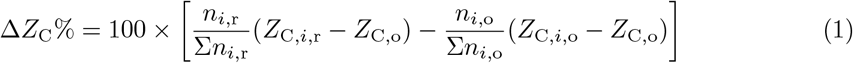

where *n*_*i*,o_ and *n*_*i*,r_ are the abundances of the *i*th taxon in the oxidized and reduced subsets, *Z*_C,*i*,o_ and *Z*_C,*i*,r_ are the carbon oxidation states of the *i*th taxon in the oxidized and reduced subsets, and *Z*_C,o_ is the carbon oxidation state of the oxidized subset. The value of *Z*_C,*i*_ was assumed to be constant for any taxon that was present in one subset but not the other. Values of Δ*Z*_C_% were visualized using the R package plot.matrix (***Klinke, 2021***).

## Author Contributions

JMD: Conceptualization, Software, Formal analysis, Writing; JT: Conceptualization, Funding acquisition.

## Data Availability Statement

All data analyzed in this study were obtained from public databases: NCBI RefSeq (***O’Leary et al., 2016***) and SRA (***Leinonen et al., 2011***); the BioProject accession numbers used to download the SRA data are listed in Table 1. The JMDplots package version 1.2.8 deposited at Zenodo (***Dick, 2021a***) has the scripts used for processing RefSeq proteins and 16S rRNA gene sequences (in the directories `extdata/refseq` and `extdata/chem16S`, respectively) and processed data files including amino acid compositions of taxa compiled from the RefSeq database `(extdata/refseq/protein_refseq.csv.xz)`, chemical metrics calculated for taxonomic groups in RefSeq `(extdata/chem16S/taxon_metrics.csv)`, and RDP Classifier results `(extdata/geo16S/RDP)`. All figures were made using R (***R Core Team, 2021***) with data files and code provided in the JMDplots package; the `geo16S.Rmd` vignette in the package runs the functions to make each of the figures and the Source Data files. The compiled vignette is available in an HTML file in the package and can also be viewed at https://chnosz.net/JMDplots/doc/geo16S.html (accessed on 2021-09-02).

## Appendix 1

**Appendix 1 Table S1.**
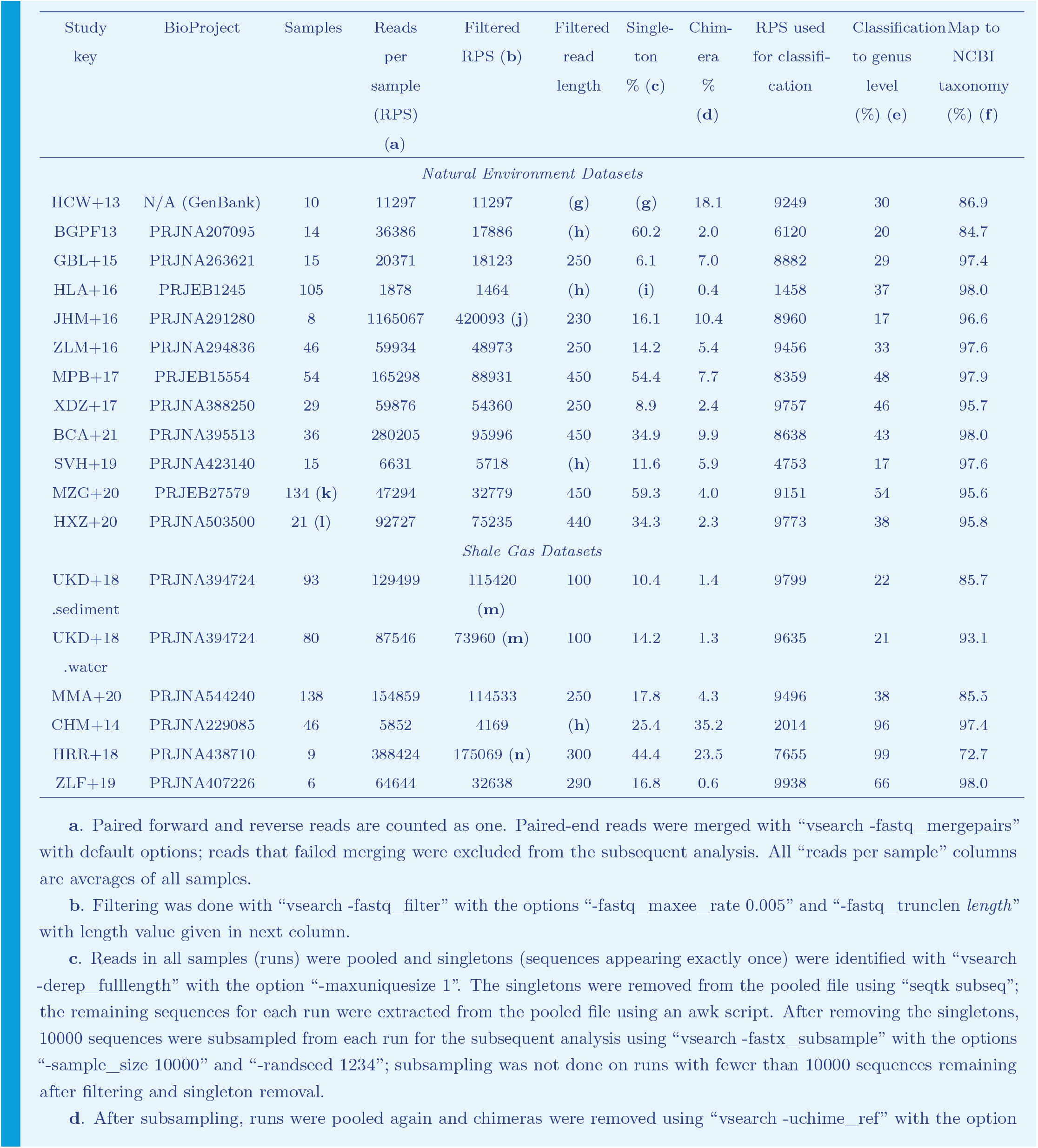
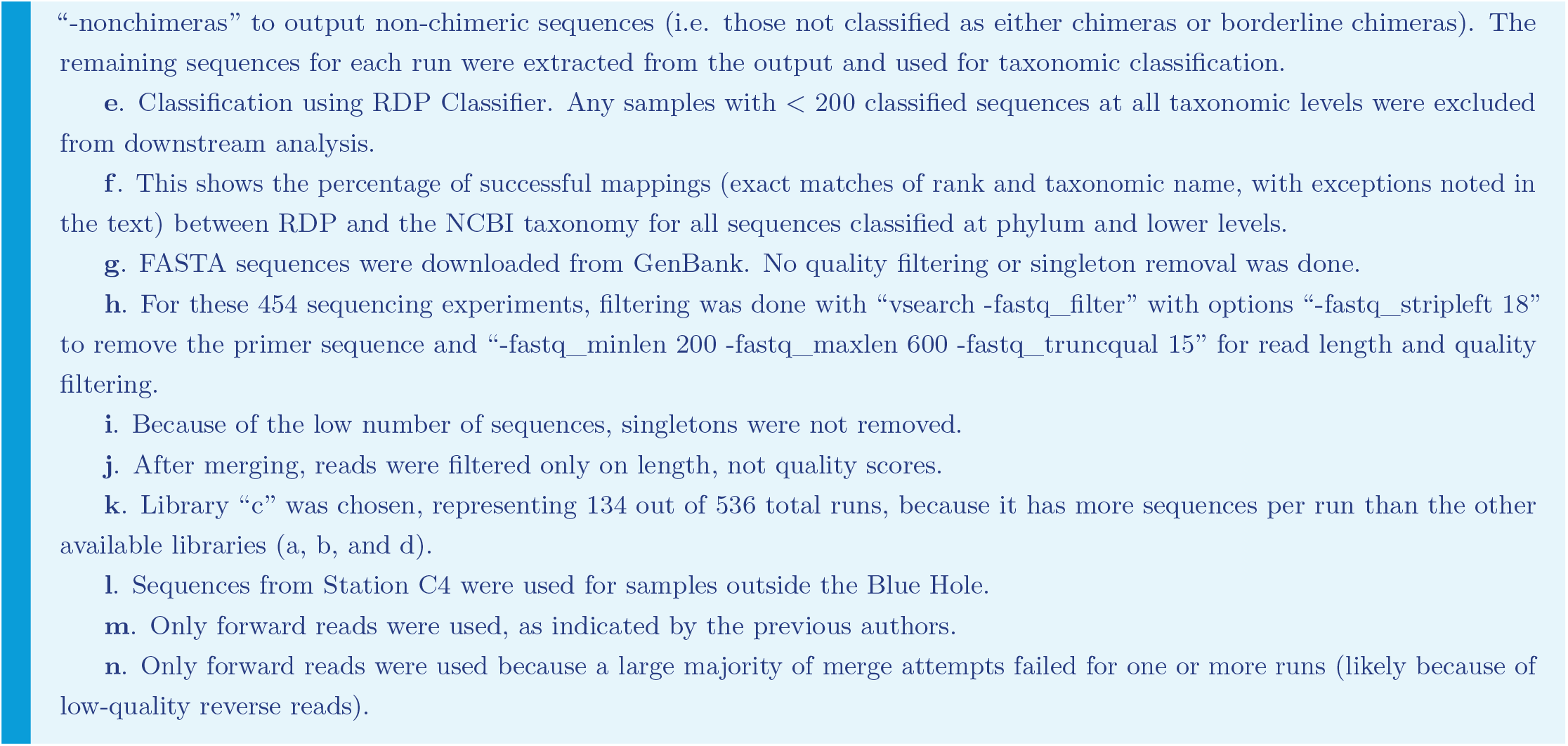
Sequence processing statistics.

**Appendix 1 Table S2.**
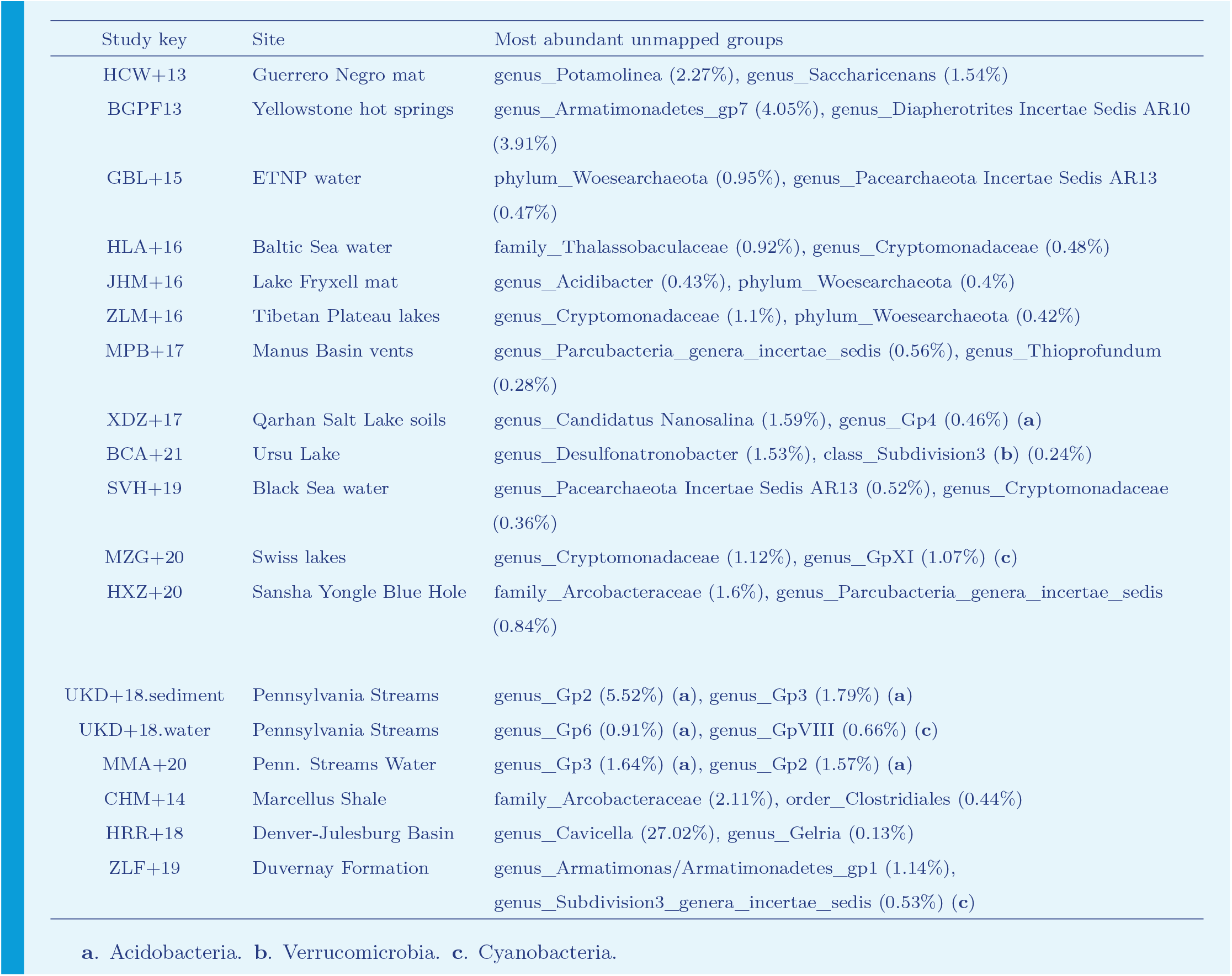
Most abundant unmapped taxonomic groups.

**Figure 6–Figure supplement 1.**
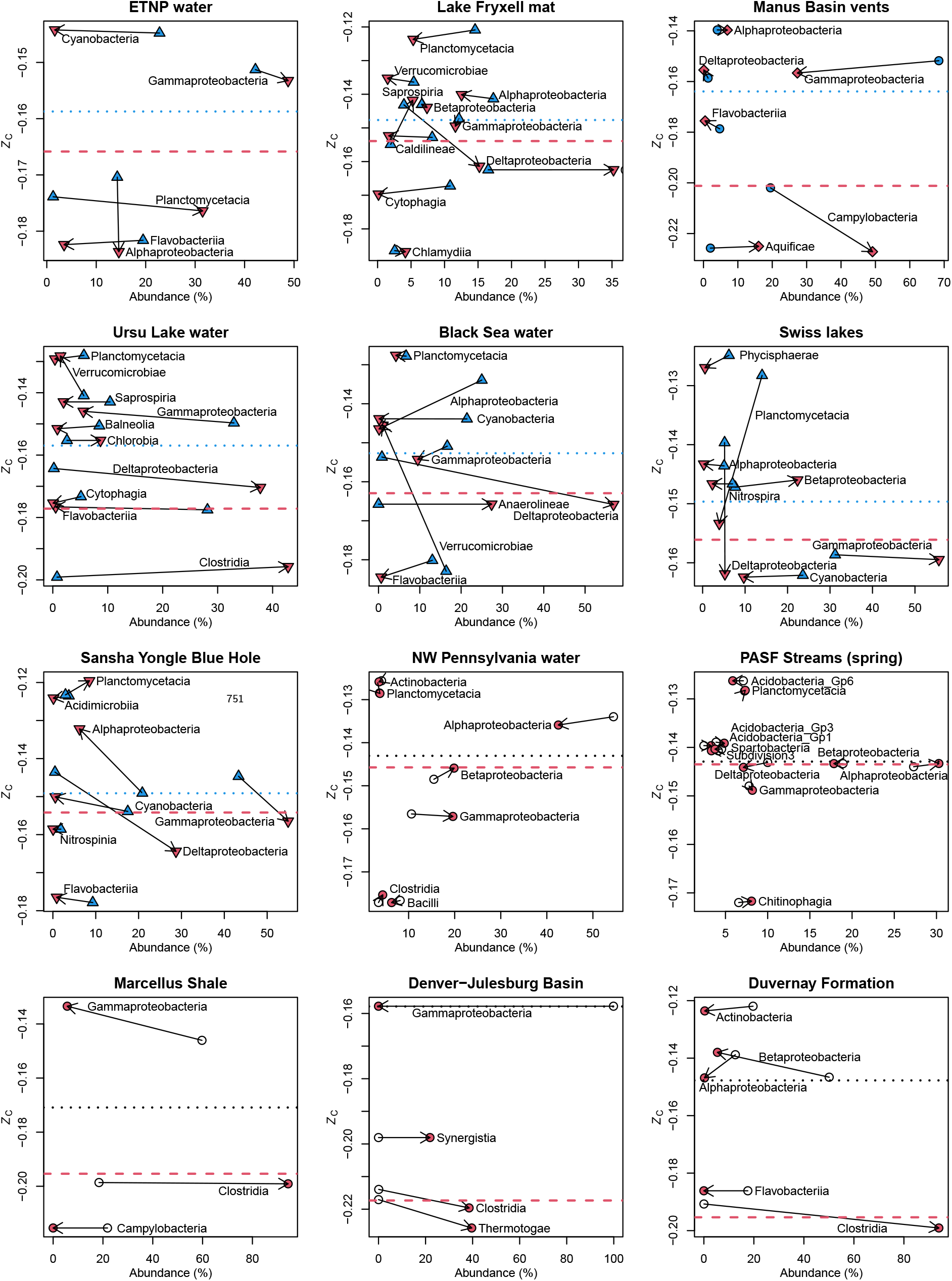
Calculations were performed as described in **Figure 4**. Arrows are drawn between subsets of samples classified as oxidized and reduced according to the descriptions in **Figure 6**

